# Systematic functional assessment of anti-phage systems in their native host

**DOI:** 10.1101/2024.12.20.629700

**Authors:** Ellie David, Clarisse Plantady, Sophiane Poissonier, Josie Elliott, Elodie Kenck, Justine Le Boulch, Arnaud Gutierrez, Anne Chevallereau

## Abstract

Bacterial resistance to bacteriophages (phages) relies on two primary strategies: preventing phage attachment and blocking post-attachment steps. These post-attachment mechanisms are mediated by diverse defence systems, including DNA-degrading systems such as Restriction-Modification (RM) and CRISPR-Cas, as along with abortive infection systems that induce cell death or dormancy. Computational analyses suggest that bacterial genomes encode multiple defence systems, which may act synergistically to enhance phage resistance. However, the regulation, interactions, and ecological roles of these systems in native hosts remain poorly understood. This study explored the role of eight predicted defence systems in the clinical isolate NILS69 of *E. coli* by testing its susceptibility to 93 phages. Infectivity and adsorption assays using mutants defective in these systems revealed that only PD-T4-3 and RM systems restricted phages able to adsorb. The RM system acted via a predicted Type IV endonuclease and was also able to limit plasmid conjugation if the plasmid was transferred from a donor strain lacking a methylase, which is the hallmark of Type I, II or III RM systems. Other defence systems showed no detectable activity, likely due to phage specificity, environmental regulation, or cofactor requirements. These findings underscore the need for further studies to investigate the regulation and ecological roles of bacterial defence systems in their native host contexts.

## Introduction

The balance between resistance and susceptibility of bacteria to their viruses (bacteriophages, or phages) is governed by two primary strategies. The first strategy consists in preventing the attachment of phages to the bacterial cell surface. This can occur through the loss, modification, or masking of phage receptors, effectively rendering the host inaccessible to infection. The second blocks post-adsorption steps in the phage lifecycle, a process mediated by specialized defence systems. In recent years, more than a hundred novel defence systems have been identified, revealing a remarkable diversity of mechanisms. These systems include those that degrade invading phage DNA, such as Restriction-Modification (RM) systems and CRISPR-Cas systems. Others inhibit phage gene expression or replication, exemplified by mechanisms like Viperins and chemical-based defences (1–3). Additionally, some defence systems rely on extreme strategies, including triggering programmed cell death or inducing dormancy, to stop phage proliferation. These mechanisms are known as abortive infection systems (4). The variety of such defences reflects the constant and dynamic evolutionary struggle between bacteria and their viral predators, as highlighted in recent literature (5). Computational analyses based on remote sequence homology searches have predicted that the bacterial defensome (i.e., the set of defence genes present in the same genome) is composed of 5-6 defence systems in average, although this number is highly variable, even between related strains (6). These findings have led to the view of the defensome as an integrated bacterial immune system, where individual defence systems are involved in a complex network of interactions. Supporting this view, synergistic associations of defence systems have been reported recently (7–12). Carrying multiple defence systems in a single genome is thought to have two main positive effects on bacterial fitness: it protects against a larger diversity of phages (11,13), as individual defence systems may have a narrow specificity and restrict only few phage genera (13), and it limits the emergence of phages that escape bacterial defences (7,11,14,15). However, we still know very little about how defence systems interact with one another, how they are regulated, and how they impact the ecology and evolution of phage-bacteria interactions. One reason for this knowledge gap is that research necessarily needed an initial effort focused on scaling up the discovery and characterisation of defence systems, which rely on novel informatic tools and synthetic biology. Consequently, most defence systems were studied from their heterologous expression in model organisms such as *Escherichia coli* and *Bacillus subtilis* (16–18). Comparatively, fewer defence systems have been studied in their original host, which hinders our understanding of the activity, regulation, fitness costs and benefits of defence genes (and their combinations) in their native condition. Recent studies have interrogated whether and how the composition of the defensome impacts the range of phages capable of infecting a host. Intuitively, one might expect a negative correlation between the number of phages infecting a host and the number and diversity of host defences. However, experimental findings have yielded contrasting results. A recent study evaluating interactions between clinical isolates of *Pseudomonas aeruginosa* and its phages found a positive correlation between the number of defence systems encoded in the isolates and their resistance to the tested phages (13). In contrast, a large-scale screen of phage-bacteria interactions in *Escherichia coli* found that, within this cohort, phage resistance was more likely determined by adsorption factors whereas a very low correlation was observed between the number of defence systems and the level of phage susceptibility (19). This suggests that, in this dataset, defence systems played only a marginal role in shaping the range of phages capable of infecting a host. Additionally, this study observed that the host range of specialist and generalist phages tended to overlap, generating an interaction matrix with a nested pattern.

In another study, the investigation of interactions between natural, longitudinally sampled, populations of *Vibrio* and their phages generated a modular interaction matrix: a specific subset of phages infected a specific subset of related hosts and thus, the host range of phages from two distinct subsets displayed very little overlap (20). In this case, the lack of interactions between phages and bacteria that belong to distinct modules is due to lack of adsorption. Conversely, the lack of interaction between phages and bacteria that belong to the same module is not explained by adsorption factors but correlates with the presence of defence systems (20). In sum, the contribution of the defensome in determining phage-bacteria ecological interactions remains unclear. In addition, the functionality and role of defence systems in their native host have been overlooked, and therefore, little is known about their levels of expression, their regulation, and their interactions with other defence systems.

In this study, we aimed to complement ecological interaction matrices by providing a genetic approach to assess the relative contribution of defence systems and adsorption factors in shaping phage host range. Specifically, we employed a systematic genetic strategy to directly evaluate the activity of predicted defence systems in their native bacterial hosts. This approach allowed us to quantify the respective contributions of individual defence systems in restricting the host range of phages or modulating their infectivity levels. By integrating these direct genetic assessments with ecological interaction data, we sought to bridge the gap between theoretical predictions and empirical observations, offering a more nuanced understanding of the interplay between phage resistance mechanisms and host susceptibility.

## Results

### Phage sensitivity of *E. coli* clinical isolate NILS69 is mainly explained by adsorption factors

To test the role of defence systems in shaping phage susceptibility, we chose the clinical *E. coli* B2 phylogroup, uropathogenic isolate NILS69 as a model system. This strain is part of a collection of *E. coli* Natural Isolates with Low Subculture passages (NILS), developed to provide the community with *E. coli* natural isolates that cover the diversity of phylogroups while remaining as close as possible to the original isolated strain (21).

We initially evaluated phage infectivity in wildtype (WT) NILS69 by screening the recently published Antonina Guelin (AG) collection, which gathers virulent dsDNA phages belonging to 19 different genera (19), through solid and liquid infection succeptibility assays (Supplementary Figure 1A,B). Ten phages out of 93 tested were able to affect the growth of NILS69 in solid infection assay amongst which six affected NILS69 growth in liquid (Supplementary Table 1).

To determine whether the infectivity of certain phages could be revealed under different culture conditions, we varied the parameters of the solid infection assay, including incubation temperature (30°C, 42°C) and salt concentration (by adding 30 mM MgSO_4_and 15 mM CaCl_2_to the growth medium), but none of these conditions changed the range of infecting phages (Supplementary Figure 1A).

We next characterized the infectivity of these 10 phages in more details. In solid medium, they form plaques with low efficiency in efficiency of plating assays (EOP ranging from 10^-5^ to 10^-1^), except for phage NIC06_P2 (Figure 1A). For some of these phages, infection on solid medium did not result in the formation of individual plaques but rather in a large lysis area. Therefore, here and for all EOP assays, we scored EOP as the highest dilution factor allowing the detection of phage lysis activity, divided by the PFU counted on the production strain. This lysis area, occurring at high phage concentration, is likely the reflection of the phage population overwhelming the host defence mechanisms. When introduced at a multiplicity of infection (MOI) of 1 in liquid medium, they show different impact on bacterial growth, which is either unaffected (536_P1, P6, P7, P9), delayed (LF73_P4, DIJ07_P1, AN24_P4) or strongly suppressed (LF73_P1, LF73_P3, NIC06_P2) (Figure 1B). The effects of phages on bacterial growth were overall consistent between liquid and solid assays: the phages with highest EOP (i.e., producing as many plaques on NILS69 as on the production strain) cause the strongest reduction of bacterial growth in liquid, whereas phages with low EOP do not affect bacterial growth. The only exception is phage LF73_P4, which has little effect in liquid infection assay despite having a relatively high EOP.

**Figure 1.**
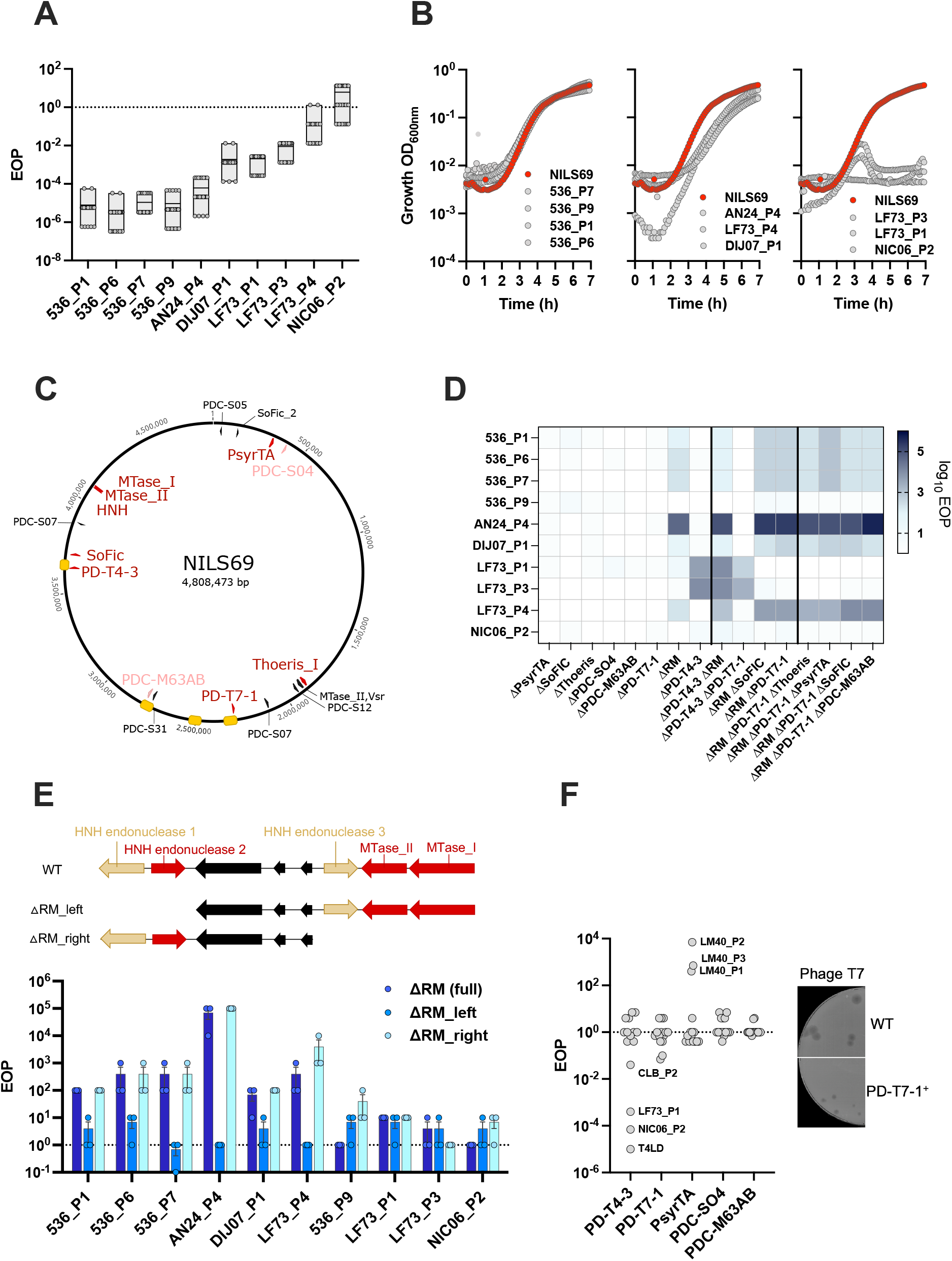
Activity of defence systems encoded in NILS69 against ten phages. **A**,**B**. Phage infectivity on NILS69 measured through Efficiency of Plating (EOP) assays, indicated as a ratio of the phage activity on NILS69 on the activity on phage production strain.(A) or liquid infection assays using a multiplicity of infection (MOI) of 1 (B). **C**. Genomic map of NILS69 indicating predicted defence systems (red), putative defence systems assessed in this study (pink) and putative defence systems not tested in this study (black). Location of prophages is indicated by yellow boxes. **D**. Phage infectivity on NIL69 defence mutants measured through EOP assays **E**. Schematic of the RM island locus in NILS69 and derivative mutants. Red arrows indicate genes that were flagged as defence genes by DefenseFinder and PADLOC, yellow arrows indicate genes that were identified as putative endonuclease by manual inspection and black arrows indicate genes encoding hypothetical proteins (upper panel). Phage infectivity was measured through EOP assays (PFU on mutant NILS69/PFU on WT NILS69) on indicated RM mutants (lower panel). **F**. Phage infectivity on K-12 expressing NILS69 defence systems in *trans* from pZS plasmids. The EOP was calculated as (PFU on K-12 pZS_defence plasmid)/(PFU on K-12 pZS_empty plasmid). Pictures of phage T7 PFU obtained on MG1655 pZS_empty or K-12 _pZS_PD-T7-1. All experiments were performed in triplicate, except for the EOP in panel A which were replicated 18 times. Boxes in panel A indicate minimal to maximal values, the line indicates the mean value and individual data points are shown. Error bars in panel E indicate standard error of mean.

Next, time-resolved adsorption assays were performed for each of the 93 phages on NILS69 to test whether the host range defined by our initial screening was due to a defect in phage adsorption. We confirmed that the 83 phages defined as non-infective by our screen failed to bind to NILS69 (Supplementary Figure 2). Therefore, the susceptibility of NILS69 to these 93 phages appears to be exclusively determined by its surface composition. Previous genome analysis indicated that NILS69 carries capsule loci while also exhibiting a rough LPS phenotype. This combination of capsular structures and the absence of full-length O-antigen may contribute to increased resistance to phage adsorption.

### Restriction Modification system and PD-T4-3 reduce the virulence of infecting phages

We next wondered whether intracellular defences could impact the infectivity levels of the 10 phages that infect NILS69. NILS69 carries 6 defence systems as predicted by PADLOC (22) and DefenseFinder(6) (Figure 1C and Supplementary Table 2), as well as 7 additional putative systems predicted by PADLOC. Individual deletion mutants of PD-T4-3, PD-T7-1, PsyrTA, SoFIC and Thoeris were constructed (Figure 1D). Regarding the predicted type II RM system, computational tools identified a HNH enconuclease associated with two methyltransferases (red arrows in Figure 1E). Manual inspection of the neighbouring region revealed two additional HNH endonucleases (referred to as HNH endonuclease 1 and HNH endonuclease 3, yellow arrows in Figure 1E) encoded in direct vicinity of the type II RM, which were not detected by prediction algorithms. A BLAST search against the REBASE database (23) indicated that HNH endonuclease 1 and HNH endonuclease 3 are similar to two putative type IV endonucleases (Osp6506McrB2P and ObaORFAP with e-values of 4E-40 and 7E-20 respectively), although these were predicted based on structural prediction and not verified experimentally. As these two endonucleases might have an effect on phages, we knocked out a large 10.6-kb region, that we refer to as RM island, encompassing the 3 HNH endonucleases as well as the two methyltransferases. Out of the seven putative defence systems predicted by PADLOC (22), we found PDC-SO4 and PDC-M63AB of particular interest as the first may be similar to a Type II Toxin Antitoxin system (according to search against TADB3.0 (24)) and the later consists In two genes that are encoded within a prophage,Ich are known to be hotspots of defence systems (6,25–27). For these reasons, we included deletion mutants of these 2 putative defence systems for downstream analyses.

The infectivities of the 10 phages were measured on each mutant through EOP (PFU on mutant NILS69/ PFU on NILS69) and liquid assays (Figure 1D and Supplementary Figure 3). Increased EOP were observed for all phages, except for NIC06_P2 and 536_P9, on mutants lacking the RM island and PD-T4-3, indicating that these defence systems protect NILS69 against these phages, while the other systems do not seem to contribute. Liquid infection assays yielded similar results, where the growth of the mutants lacking the RM island and PD-T4-3 was effectively suppressed by phages. While 536_P9 did not formed more plaques on any mutant in solid assays, this phage could effectively suppress the growth of the RM island mutant in liquid, indicating that RM is also active against this phage (Supplementary Figure 3).

We next wondered if the phages that still induce weak lysis of NILS69 despite being restricted by the RM system carry escape mutations or epigenetic modifications. To test this, we collected phages by scraping from the lysis area formed at high phage concentration by phages 536_P6 and 536_P7 growing on NILS69 and used them to reinfect both NILS69 and the production strain 536. We predicted that, if the recovered phages were escaping NILS69 defences because of mutations or genome modifications, they would lyse NILS69 and 536 with similar efficacy. However, the recovered phages formed plaques on 536 but not on NILS69, indicating that they were not carrying escape mutations or modifications (Supplementary Figure 4), but were rather the product of a large phage population overwhelming the defence capacity of NILS69.

The RM island and PD-T4-3 target a non-overlapping set of phages, with the RM island restricting 7 phages (536_P1, 536_P6, 536_P7, 536_P9, AN24_P4, DIJ07_P1, LF73_P4) and PD-T4-3 specifically inhibiting LF73_P1 and LF73_P3. To investigate potential negative or positive interactions between defence systems, we generated four double mutants and four triple mutants and tested their phage sensitivity using solid assays. Among these mutants, only the double mutant lacking both PD-T4-3 and the RM island exhibited increased sensitivity to phages, which was comparable to the additive effect observed in the corresponding single mutants (Figure 1D). We used these mutants lacking multiple defence systems to screen the AG phage collection again and confirmed that the lack of infectivity of the 83 phages is not due to defence activity (Supplementary Figure 1A).

Altogether, our data indicate that the RM island and PD-T4-3 are the only active defence systems in NILS69 under these conditions and against this collection of phages, and that their combined activity provides an additive protection against a broader range of phages.

### Experimental validation of the activity of an HNH endonuclease

Next, we aimed to identify which endonuclease within the RM island mediates phage protection, hypothesizing that the type II RM system detected by DefenseFinder and PADLOC (HNH endonuclease 2), which shows 100% identity to several restriction enzymes in REBASE, is likely responsible for this phenotype. We constructed two mutants with deletions in different portions of the RM island and measured phage EOP (Figure 1E). Only the mutants lacking HNH endonuclease 3 (ΔRM_right) exhibited a phage sensitivity profile comparable to the mutant lacking the entire RM island. These findings indicate that the type II endonuclease detected by DefenseFinder and PADLOC is not responsible for phage protection. A sequence homology search using BLASTP against the Standard database revealed that HNH endonuclease 3 has 100% identity to an HNH endonuclease found in other *E. coli* isolates (Uniprot: A0A3K0QCZ9). A structure-based homology search using AlphaFold3 (28) and FoldSeek (29) identified a match to the Type IV methyl-directed restriction enzyme EcoKMcrA encoded by *E. coli* K-12 MG1655 (E-value: 2.48E-04, run in Nov. 2024). Our data indicate that the previously untested HNH endonuclease 3 has an antiphage activity and shares structural features with a Type IV restriction enzyme (Figure 1E, Supplementary Figure 5).

### Evaluation of the activity of NILS69 defence systems in the *E. coli* laboratory K-12

The apparent lack of activity of most of predicted defence systems might be linked to their target specificity. In other words, they might be active against other phages than those we tested. To explore this possibility, we attempted to cloned each individual defence system under its native upstream regulatory sequence into a low copy plasmid and subsequently transformed the constructs in *E. coli* K-12. Despite repeated attempts, we were unable to clone Thoeris and the RM island. The K-12 strain has a distinct phage sensitivity profile compared to NILS69, being sensitive to 15 of the 93 phages tested (Supplementary Figure 1A). Comparing the infectivity of the 15 phages on K-12 and on clones complemented with individual defence systems confirmed that PD-T4-3 is active and protects against 4 phages (CLB_P2, LF73_P1, NIC06_P2 and T4LD) which all belong to the *Tevenvirinae* subfamily (Figure 1F). Previous work indicated that PD-T4-3 targets phages from the *Tequatrovirus* genus (T2, T4 and T6)(18). Our data suggest that the specificity of PD-T4-3 is not limited to *Tequatrovirus* as it targets phage from *Dhakavirus* genus (CLB_P2) but PD-T4-3 likely does not target all genera from the *Tevenvirinae* subfamily since phages from *Mosigvirus* (LF82_P8) do not seem to be affected. Interestingly, phage NIC06_P2, which is unaffected by PD-T4-3 in NILS69, becomes sensitive to this defence system when overexpressed in K-12. This suggests that NIC06_P2 may possess a counter-defence strategy that is no longer effective under this condition of defence overexpression (Figure 1F). This assay also revealed that PD-T7-1 protects against the phage PDP351_P2 (*Teseptimavirus*) and has a partial protective activity as it reduces the plaque size of phage T7. None of the other defence system had a detectable protective activity in these conditions (Figure 1F). Surprisingly, the EOP of phages LM40_P1, LM40_P2 and LM40_P3 were higher when the strain expressed PsyrTA. Because we also noticed that this strain had a growth defect, we tested whehter the effect observed on LM40 phages was due to bacterial density. We show that phages LM40_P1, LM40_P2 and LM40_P3 have higher infectivity on low density bacterial lawns, and that the effect we see is not spectific to PsyrTA (Supplementary Figure 6).

### Restriction modification system of NILS69 limits plasmid transmission

We hypothesized that some defence systems in NILS69 may target other types of mobile genetic elements in addition to phages. Therefore, we assessed their capacity to limit plasmid transmission. We assessed the efficiency of transfer of a pRCS30, a 157kb IncC plasmid carrying multiple antibiotic resistance genes including an extended-spectrum β-lactamase gene CTX-M14(30), from its original host strain *E. coli* 513 to NILS69 and its derivative mutants.

The conjugation efficiency, measured as the ratio of transconjugants to recipients, was approximately 10^−7^ in NILS69, peaking at 1 hour. Similar efficiency was observed in all mutants except for the ΔRM strain, which exhibited a 100-fold increase (∼10^−5^), suggesting that the RM island restricts plasmid horizontal transmission. (Figure 2A). To test which endonuclease from the island was mediating this effect, we measured conjugation efficiency in the ΔRM_right and ΔRM_left mutant backgrounds and found that the same HNH endonuclease 3 was responsible for protection against both phages and plasmids (Figure 2B).

**Figure 2.**
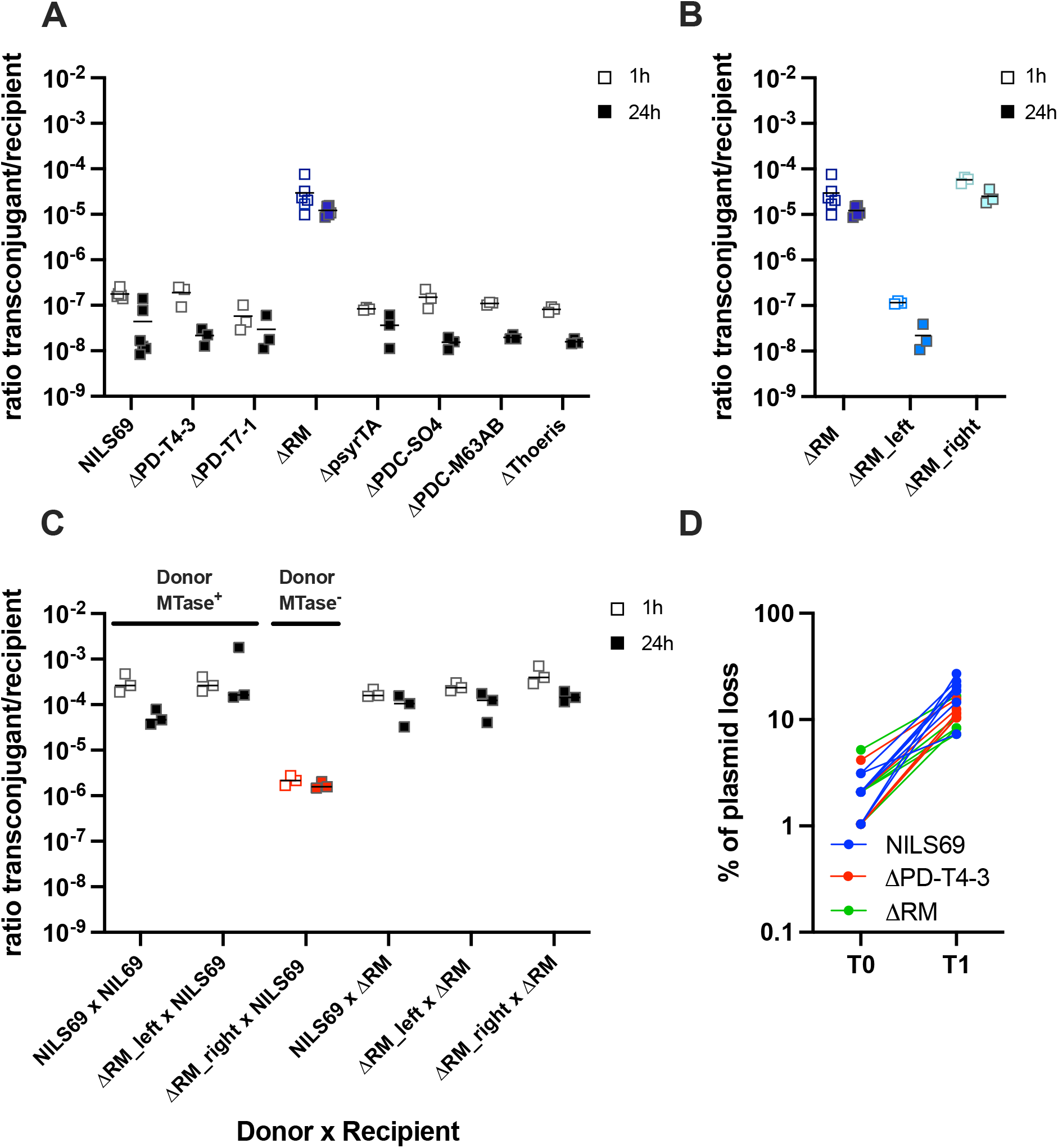
NILS69 RM system limits plasmid horizontal transmission. **A**,**B**. Conjugation efficiency of the 157kb-plasmid pRCS30 into NILS69 and defence mutant derivatives. The plasmid was transferred from its native host (*E. coli* 513). Conjugation efficiency is indicated as the ratio of transconjugant Colony Forming Units (CFU) on recipient CFU after 1h or 24h of contact between the donor and the recipient strains. **C**. Conjugation efficiency of pRCS30 from NILS69 to NILS69. **D**. Vertical transmission of pZS_GFP plasmid in NILS69 and the ΔRM and ΔPD-T4-3 mutants. All experiments were perfomed in triplicates, except in Panel A, conjugation in NILS69 and ΔRM were performed six times. Individual data points are indicated, lines indicate mean values.

Because this endonuclease shares some similarity to a putative Type IV endonuclease, we aimed to verify if it was indeed targeting methylated invading DNA or not. To test this, we recovered transconjugants in NILS69, ΔRM_right and ΔRM_left backgrounds and measured the efficiency of a secondary transfer of pRCS30 in NILS69 and ΔRM island mutants. Plasmids from the donor strains NILS69 and ΔRM_left should be methylated, as these strains encode the two methyltransferases in the RM island, while plasmids from the donor ΔRM_right strain are expected to be unmethylated (Figure 1E). First, we observed that the transfer of pRCS30 into NILS69 is 1000-fold higher when the plasmid had previously replicated in the same background compared to its original host *E. coli* 513 (Figure 2C). Second, we confirmed that this increased conjugation efficiency is due to the presence of methyltransferases in the donor strains, as their absence prevents this increase in transfer efficiency (Figure 2C). Our data therefore indicate that NILS69 encodes an HNH endonuclease that restricts plasmid and phage infection and which activity is likely inhibited by methylation.

Finally, we tested whether plasmid vertical transmission could be affected by the two defence systems that are active in NILS69: PD-T4-3 and RM. To this aim we transformed the low copy plasmid pZS into NILS69 WT, ΔRM or ΔPD-T4-3. Plasmid-carrying bacterial clones were initially selected in the presence of antibiotics and subsequently transferred on non-selective media to measure the percentage of plasmid loss in each background (Figure 2D). While a fraction of the population lost the plasmid, this did not depend of the bacterial genetic background, therefore RM and PD-T4-3 had no effect on the vertical transmission of this plasmid.

## Discussion

The role of defence systems in shaping phage-bacteria interactions is still unclear. In particular, the benefits provided by the accumulation of defence systems in a single genome remain an open question. In this study we used a direct genetic approach to investigate the impact of defence systems on the vulnerability of the *E. coli* clinical isolate NILS69 to a diverse collection of phages. Confirming a previous study, we experimentally show that the inability of phage to produce a successful infection is mainly explained by the lack of adsorption. Out of the six predicted defence systems, only two have a measurable impact on the infectivity of the phages that can bind the host; Restriction Modification and PD-T4-3. In addition, we tested two putative systems (i.e., never previously verified experimentally) predicted by PADLOC(22,31), namely PDC-M63AB and PDC-S04, but could not find evidence of defence activity in our model system. These findings raise questions about the activity of other predicted defence systems. We can foresee several reasons for the apparent inactivity of these systems. First, the limited diversity of the phage collection, which consists of virulent, double-stranded DNA phages, may introduce bias. Previous work has showed that certain defence systems are specific and target a narrow range of phages (13,18) and it may be that we have just not tested the phages that are sensitive to these defence systems.

Curiously, phage NIC06_P2 can be targeted by PD-T4-3 when expressed in *trans in E. coli* K-12, but is not targeted in the native context of NILS69.

Therefore, a second possibility is that some of these defence systems may be strongly regulated and expressed in different environmental conditions or may require additional cofactor to act. Although some regulators have been identified (11,32,33), whether defence systems are expressed and differentially regulated depending on the environment remains an unexplored area. Future studies aiming at measuring the impact of the environmental context on defence systems activity and their consequences on phage activity will be key to advance our understanding on the ecological impact of defence systems.A third explanation to the apparent inactivity of defence systems is given by phage-encoded counter-defence proteins. Many counter-defences have been identified recently in phages or plasmids and it clearly appears that many more remains to be uncovered (34,35). A recent effort was made to catalogue and detect all known counter-defence proteins from genomes (36). In this study, we observed that the phage NIC06_P2, which has the highest EOP on NILS69 despite an apparent low adsorption efficiency (Figure 1A, Supplementary Figure 2), is unaffected by the defence systems tested since it lyses NILS69 as efficiently as all the derivative mutants. Interestingly, this phage encodes a predicted anti-restriction nuclease (Arn) protein, which, in phage T4, inhibits the modification-dependent endonuclease McrBC (Supplementary Table 1). Phage LF73_P4 also encodes a predicted anti-RM protein, Dam, which may partially protect the phage against the RM island. In addition, NIC06_P2 is unaffected by the defence system PD-T4-3, while closely related phages are (LF73_P1 and LF73_P3). PD-T4-3 is a single gene system that encodes a GIY-YIG nuclease domain but its mechanism of action is unknown. Our data suggest that NIC06_P2 may escape PD-T4-3 in NILS69, but not in K-12 where this system is likely overexpressed (due to plasmid-based expression or to a lack of repression that might be at play in the native host).

Interrogating the effect of defence systems on other mobile genetic elements, we found that an HNH endonuclease effectively limits both phage and plasmid horizontal transmission. Interestingly, this endonuclease shares structural similarity to a Type IV methylcytosine-targeting restriction enzyme but our data indicate that the methyltransferase that are encoded directly downstream on the opposite strand limit the endonuclease activity, suggesting a Type I, II or III mode of action. This association of an HNH endonuclease with two methyltransferases might be conserved as our preliminary analysis detected this 3-gene association in *E. coli, Enterobacter bugandensis, Klebsiella pneumoniae* and *Vibrio alginolyticus*. Further molecular investigation of this endonuclease and associated methyltransferases will help to clarify their mechanism of action.

In conclusion, we conducted a comprehensive functional assessment of defence systems in a clinical *E. coli* isolate. Our study indicates that, for this strain and in the conditions tested, the main phage barrier is the bacterial envelope, while defence systems play a more marginal role and further highlights the need to study defence systems in their native hosts, in variable environmental conditions to improve our understanding of their roles in the ecology of phage-bacteria interactions.

## Material and Method

### Bacterial strains, phages, plasmids and growth conditiond

Bacterial strains, phages, plasmids and primers used in this study are listed in Supplementary Table 3. The medium used for all experiments was Lysogenic Broth (LB) prepared according to the Luria formulation. Unless otherwise specified, bacterial cultures were inoculated in LB from frozen stocks and propagated at 37°C with agitation. When needed, antibiotics were added at the following concentrations: ampicillin 100 μg/mL, kanamycin 100 μg/mL, chloramphenicol 30 μg/mL, streptomycin 30 μg/mL, and rifampicin 100 μg/mL. Solid or soft media were prepared by adding 15 g/L or 7 g/L of agar, respectively.

### Phage efficiency scoring

Phage efficiency of plating (EOP) against NILS69 and K-12-MG1655 derivative was tested by spotting on soft agar plate 3μL of 10-fold serially diluted phage suspensions. Soft agar was inoculated to a final concentration of ≈10-^7^ CFU/mL from exponential phase growing cultures and then incubated overnight at 37°C. EOP was calculated by dividing the Plaque-Forming Units (PFU) or the highest dilution factor allowing the detection of phage lysis activity obtained on NILS69 by the number of PFU obtained on the phage production strain.. All the bacterial growth kinetic have been done in microtiter plate using “Tecan Infinite M Nano” microplate reader. Wells were inoculated with 10^5^ CFU/mL from overnight culture with or without phages at a multiplicity of infection of 1. OD600nm readings were taken every 5 minutes over an 9h incubation period at 37°C with constant agitation

The screening of E. coli NILS69 against the Antonia Guelin phage collection was performed in microtiter plate using 200 μL culture inoculated with 10^5^ CFU/mL from an overnight culture and phages at a 100x dilution from stocks. OD readings were taken every 5 minutes over a 9h incubation period at 37°C with constant agitation.

### Adsorption assays

Adsorption assays on NILS69 were conducted according to a previously described protocol (37). Overnight bacterial cultures incubated at 37°C were transferred into a deep-well plate. Phages were introduced at MOI 0.001 in a final volume of 1mL, with each well representing a unique phage/bacteria combination. Adsorption was performed at 37°C for 30 minutes without agitation. To monitor adsorption kinetics, 150 μL samples were collected at 10, 20, and 30-minute intervals, treated with 50 μL of chloroform, and the supernatants were titrated using a spot assay. This included a negative control with phages only (no bacteria) and a positive control with phages and their reference bacterial strain, both of which were also titrated.

### Deletion mutants of NILS69

Deletion mutants of the defence systems in NILS69 were constructed using site-specific recombination mediated by the **λ**-Red system, as described by Datsenko and Wanner (38). The primers listed in Supplementary Table 3 were used to amplify resistance cassettes via PCR, generating linear DNA fragments containing homology regions for recombination. This linear DNA was introduced into bacteria expressing the **λ**-Red system by electroporation, and recombinant bacteria were selected on the appropriate antibiotic-containing media. When necessary, the resistance marker was removed using FRT recombination mediated by the expression of the Flp recombinase, as described in the same study (38). All generated mutants were sequenced (Illumina, NextSeq) to confirm the deletion. The reference genome of NILS69 can be accessed on NCBI (accession: DABQWS000000000.1)

### Subcloning of defence systems and expression in E. coli K-12 MG1655

The low-copy number plasmid pZS_Cat was used to subclone NILS69 defence systems and expressed them in the lab strain K-12 MG1655. Defence systems were amplified using the PCR primer listed in Supplementary Table 3, and were cloned in the pZS_Cat vector using commonly used restriction digestion and ligation. Constructions were verified by Sanger sequencing. The primers were designed in order to include a minimum of 200 nucleotide upstream the start codon, in order to include any native promotor and regulatory sequences. Only 5 out 8 defence systems could be cloned using this method.

### Plasmid Vertical Transmission

The low-copy-number plasmid pZS_sfGFP was used to assess the plasmid vertical transmission ability of NILS69 and its derivatives. This vector, derived from the pZ vector series described by Lutz and Bujard (39) includes a low-copy-number replication origin, a kanamycin resistance gene, and allows for the constitutive expression of superfolder-GFP (sfGFP) under the control of an unregulated tetpromoter.

Vertical transmission of pZS_sfGFP was monitored as follows: overnight cultures were diluted to approximately 1–10 bacteria in media containing kanamycin and incubated for 24 hours (T1). The cultures were then diluted again to approximately 1–10 bacteria in media without antibiotic selection and incubated for an additional 24 hours (T2). At T1 and T2, the colony-forming units (CFU) were estimated by plating the cultures on solid media without selection. To estimate the proportion of bacteria retaining the plasmid, ≈100 CFUs were replicated onto plates containing kanamycin.

### Plasmid Horizontal Acquisition

The IncC family conjugative plasmid pRCS30 was used to assess the ability of NILS69 and its derivatives to acquire horizontally transferred DNA. This 157 kb plasmid, isolated from a clinical *E. coli* strain, carries various antibiotic resistance genes, including aph(6)-Id, which confers resistance to streptomycin. Since the spontaneous mutation frequency leading to streptomycin resistance in NILS69 is below 10^−9^, resistance to streptomycin was used as a marker to track the transfer of the pRCS30 plasmid. To evaluate plasmid acquisition, recipient NILS69 and its derivatives, grown to an OD_600_ of 0.2, were mixed at a 1:1 ratio with the pRCS30 donor strain and incubated statically. All recipient cells carried the stable high-copy-number plasmid pZS_sfGFP-kanR, which constitutively expressed GFP and conferred resistance to kanamycin. After 1 hour and 24 hours, the mixed cultures were vortexed vigorously for 1 minute, and bacteria were plated on LB agar containing kanamycin to estimate the total count of recipient cells (kanR). Additionally, the cultures were plated on LB agar containing both streptomycin and kanamycin to determine the number of recipient cells that acquired the pRCS30 plasmid (kanR strepR). Conjugation efficiency was calculated as the ratio of kanR strepR / kanR.

Conjugation assays across NILS derivatives were performed by first selecting spontaneous mutants resistant to rifampicin (rifR), by plating the relevant strain on LB agar containing 100 μg/mL of rifampicin. These mutants were subsequently used as recipients during conjugation assays with NILS69 derivatives carrying the pRCS30 plasmid. Here, the conjugation efficiency was calculated as the ratio of rifR strepR / rifR.

### Bioinformatic and data analysis tools

Defence systems from NILS69 genome were predicted using webservers of DefenseFinder (6,40,41) (https://defensefinder.mdmlab.fr/) and PADLOC (22,31 (https://padloc.otago.ac.nz/padloc/). Toxin Antitoxin systems were predicted with TADB 3.0 (https://bioinfo-mml.sjtu.edu.cn/TADB3/) Search for sequence and structural homologies of the HNH endonucleases were done using BLAST(42) (https://blast.ncbi.nlm.nih.gov/Blast.cgi?PAGE=Proteins) against the Rebase database(23) (https://rebase.neb.com/rebase/rebase.html), AlphaFold3(28) (https://alphafold.ebi.ac.uk/) and FoldSeek(29) (https://search.foldseek.com/search).

Graphs and data analyses were generated with Prism (GraphPad).

## Data accessibility

All raw data generated during these study have been deposited on https://github.com/MMSB-MEEP-lab

## Acknowledgments

We acknowledge Dr Luce Landraud for proving the plasmid pRCS30 and Prof. Erick Denamur for sharing the strain NILS69. The Antonina Guelin phage collection and associated production strains were a kind gift from Dr. Laurent Debarbieux and Dr. Aude Bernheim. AG is supported by the following fundings ANR-21-CE35-0003, Emergence en recherche 2020 de l’Idex Université Paris Cité RM99J20IDXA8 and Emergence ville de Paris 2020-DAE78-EMERGENCE. AC is supported by the ATIP-Avenir program and IdEx Université Paris Cité ANR 18 IDEX 0001.

## Electronic Supplementary material

**Supplementary Figure 1.**
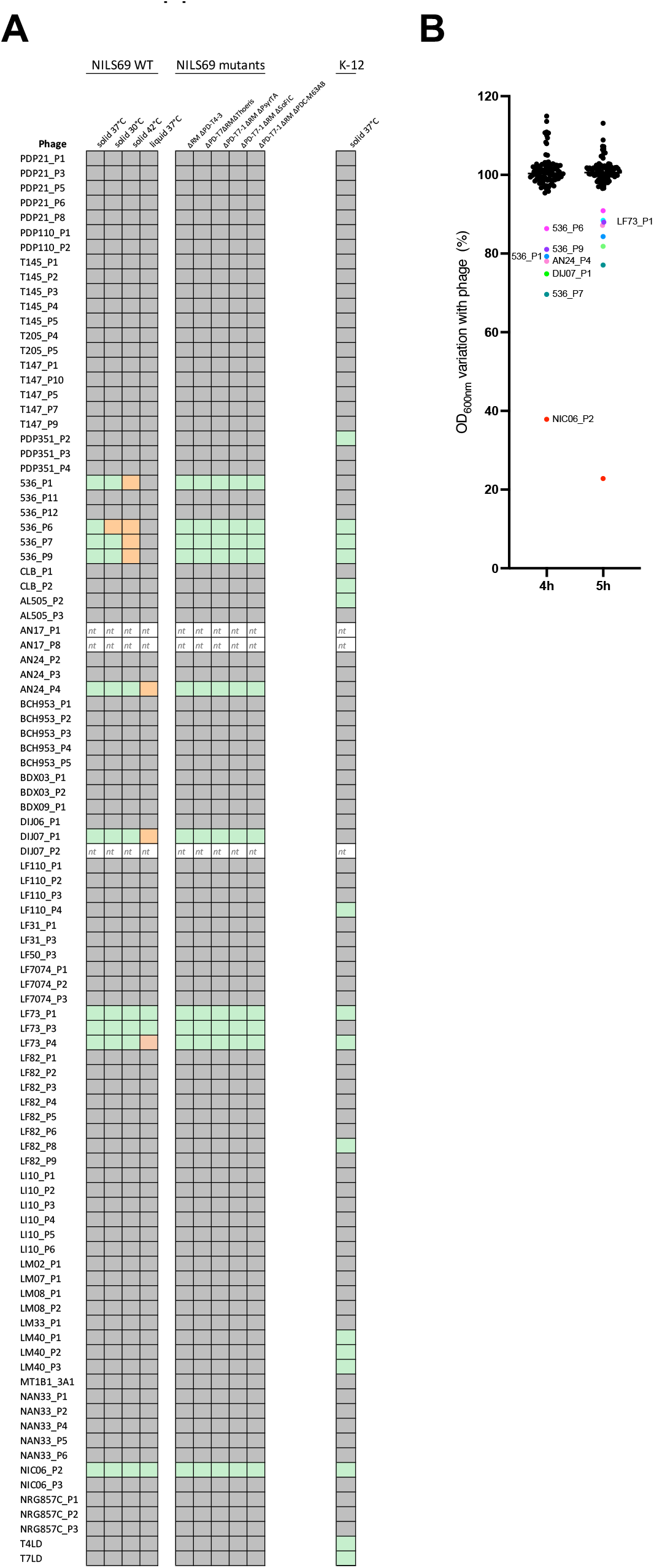
Screening of the Antonina Guelin phage collection on NILS69 and K-12 in solid (A) and liquid (B) infection assays. “nt” stands for “not tested”, the corresponding phages could not be amplified on their production strain. Green and orange squares indicate detectable phage infection (i.e. plaques). Orange square indicate cases where only faint lysis could be observed. Grey indicates no lysis

**Supplementary Figure 2.**
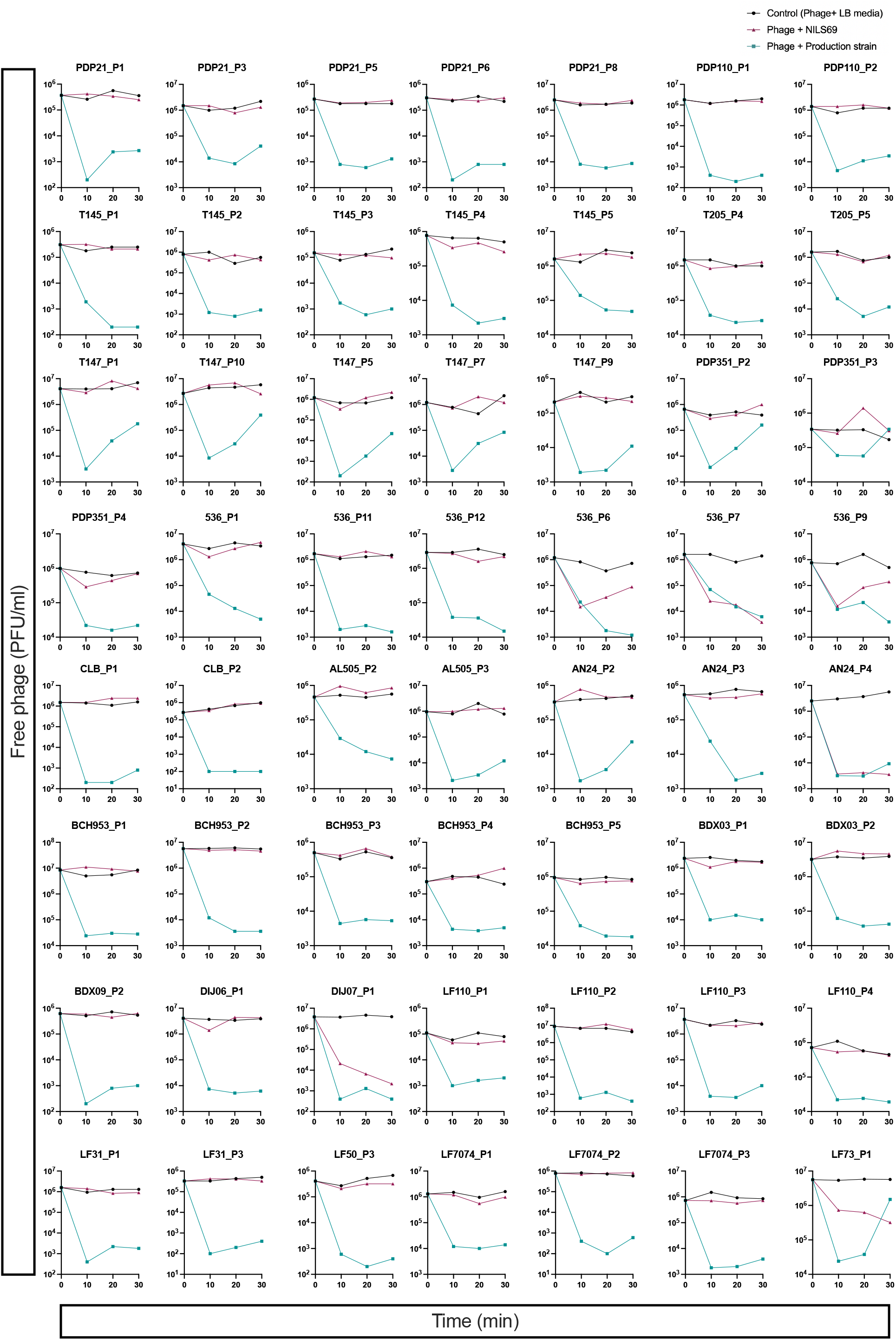
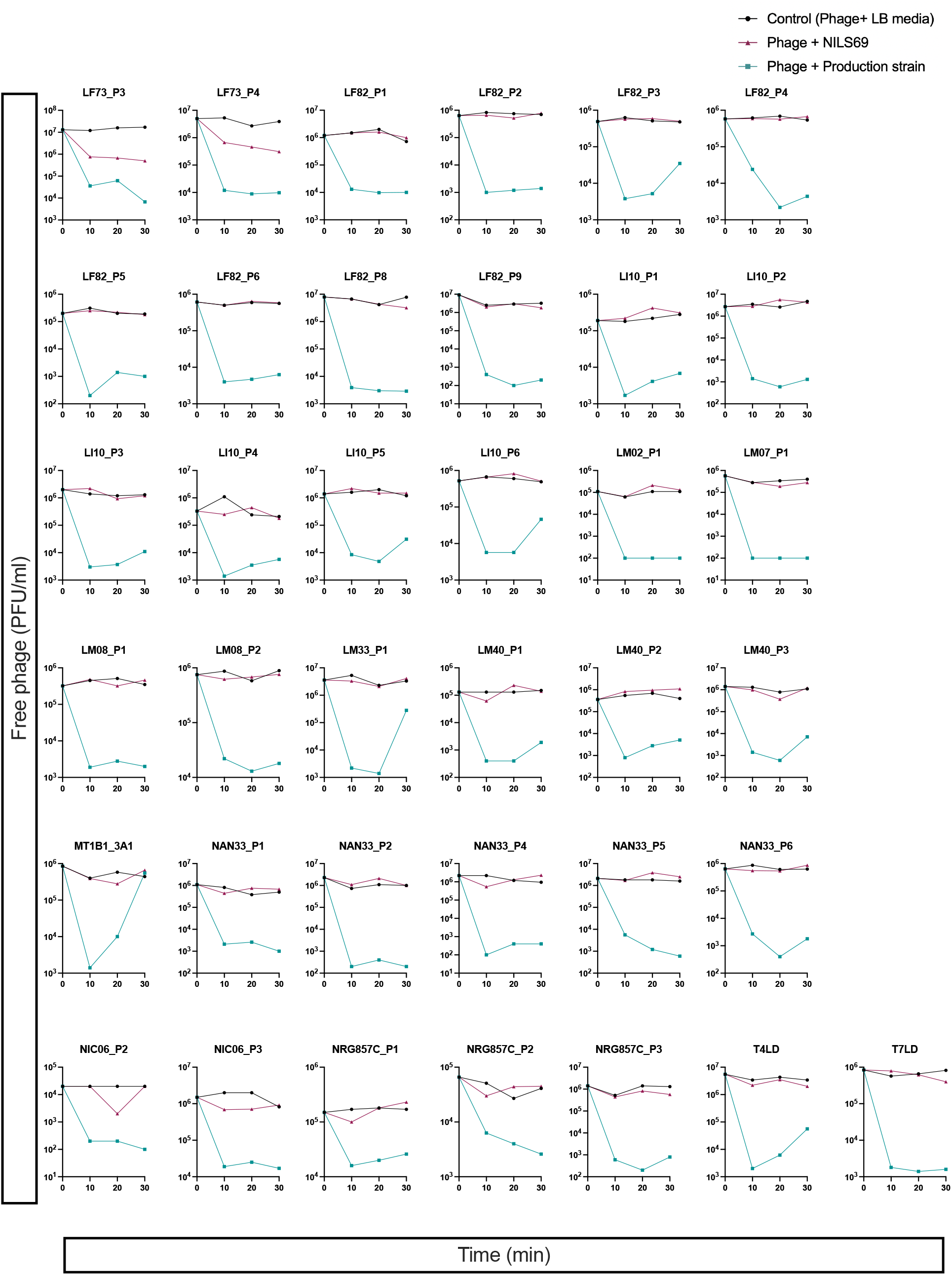
Adorption assays of 93 phages from the Antonina Guelin on NILS69 and on their control strains (phage production strains)

**Supplementary Figure 3.**
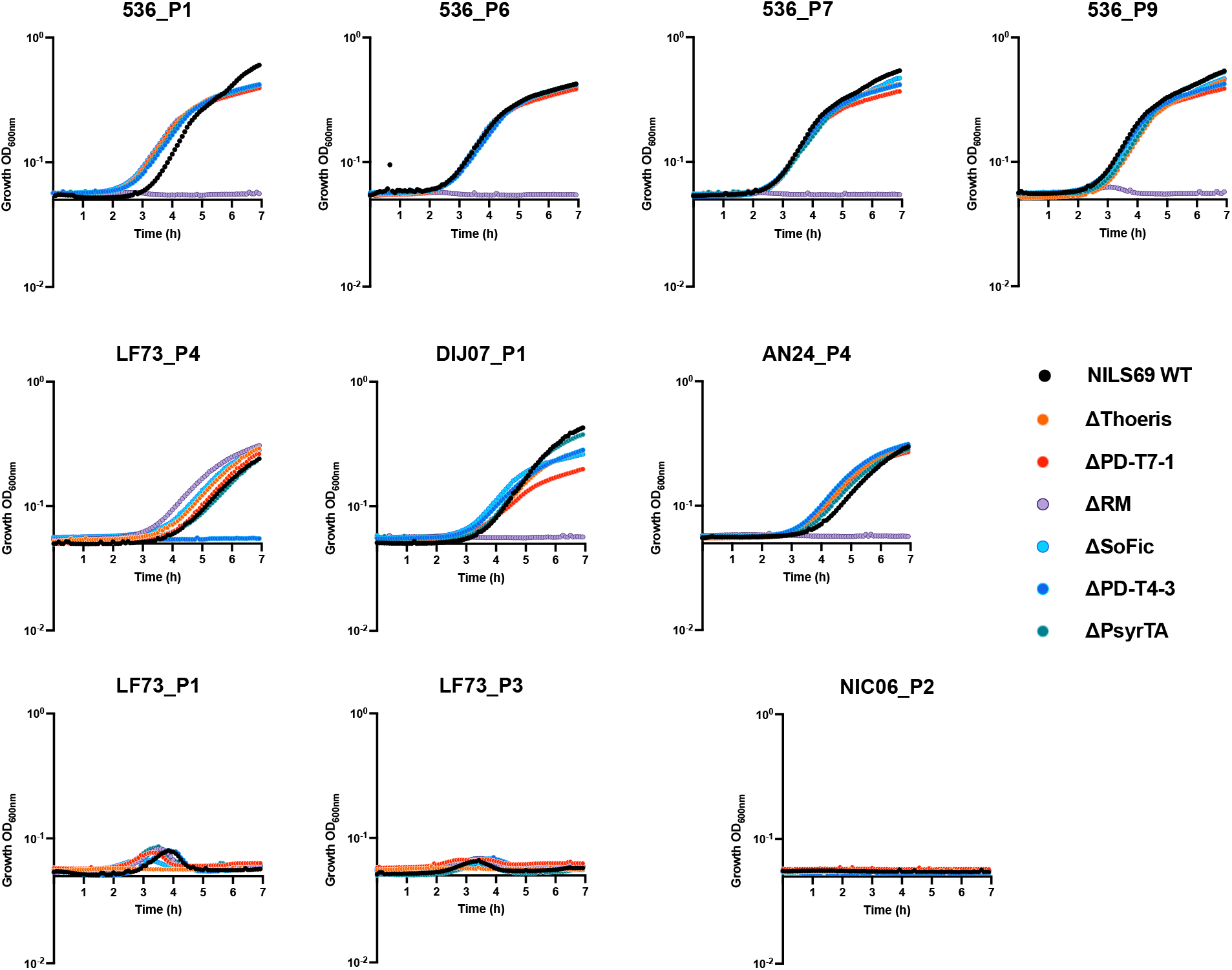
Growth kinetic of NILS69 and mutant dertivative in the presence of the infective phages at a MOI of 1.

**Supplementary Figure 4.**
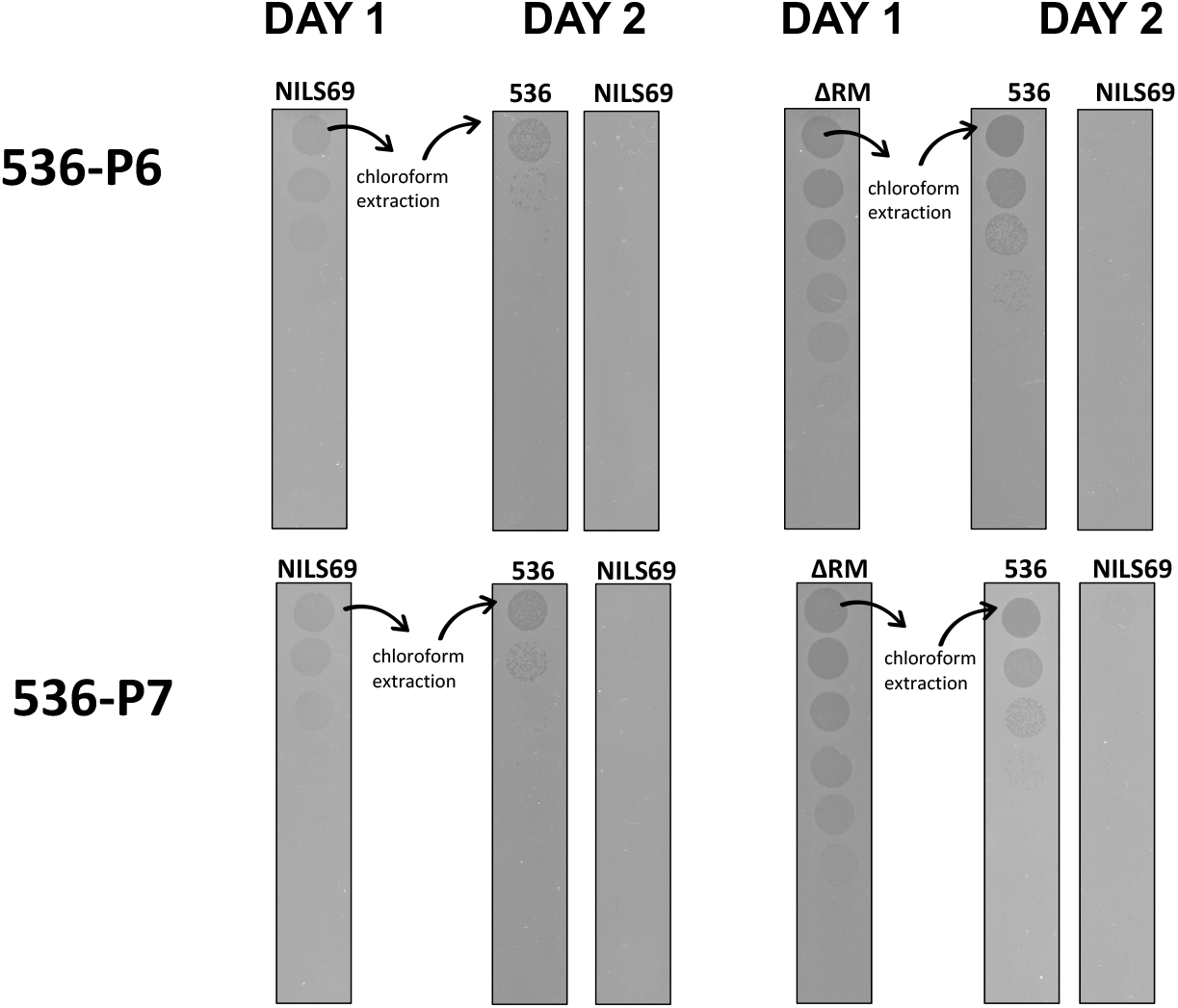
Assessment of escape phage mutants. Phages 536_P6 and 536_P7 lysing NILS69 or NILS69ΔRM were harvested from the plate and purified through chloroform extraction (Day 1). These phages were subjected to 10-fold serial dilutions and spotted on lawn of strain 536, NILS69 or NILS69ΔRM (Day 2).

**Supplementary Figure 5:**
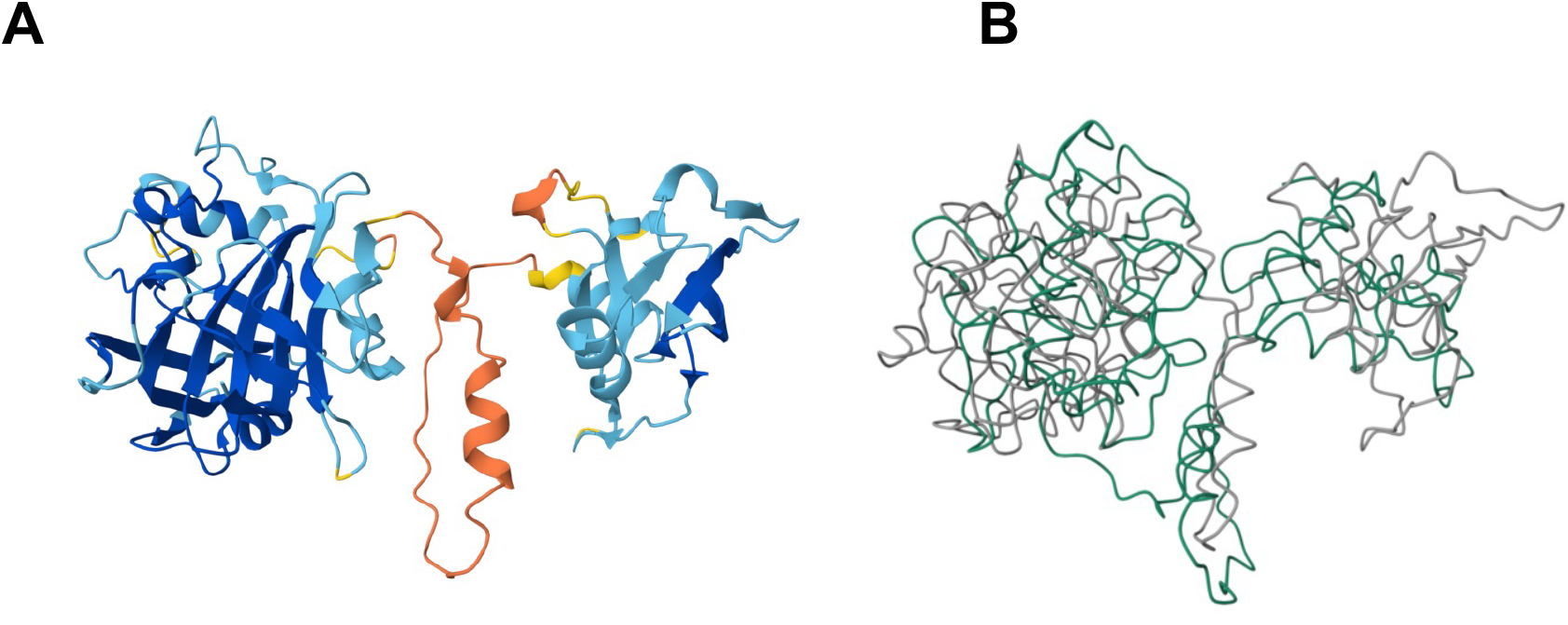
Predicted structure of HNH endonuclease 3. A. HNH endonuclease 3 encoded in NILS69 has 100% identity with the amino acid sequence of a HNH endonuclease (Uniprot A0A3K0QCZ9), which has a predicted structure in AlphaFold (average pLDDT score of 80,56) B. Structural alignment of NILS69_HNH endonuclease and K-12 MG1655_EcoKMrcA (e-value 2.48e-4; sequence identity 20.6%)

**Supplementary Figure 6:**
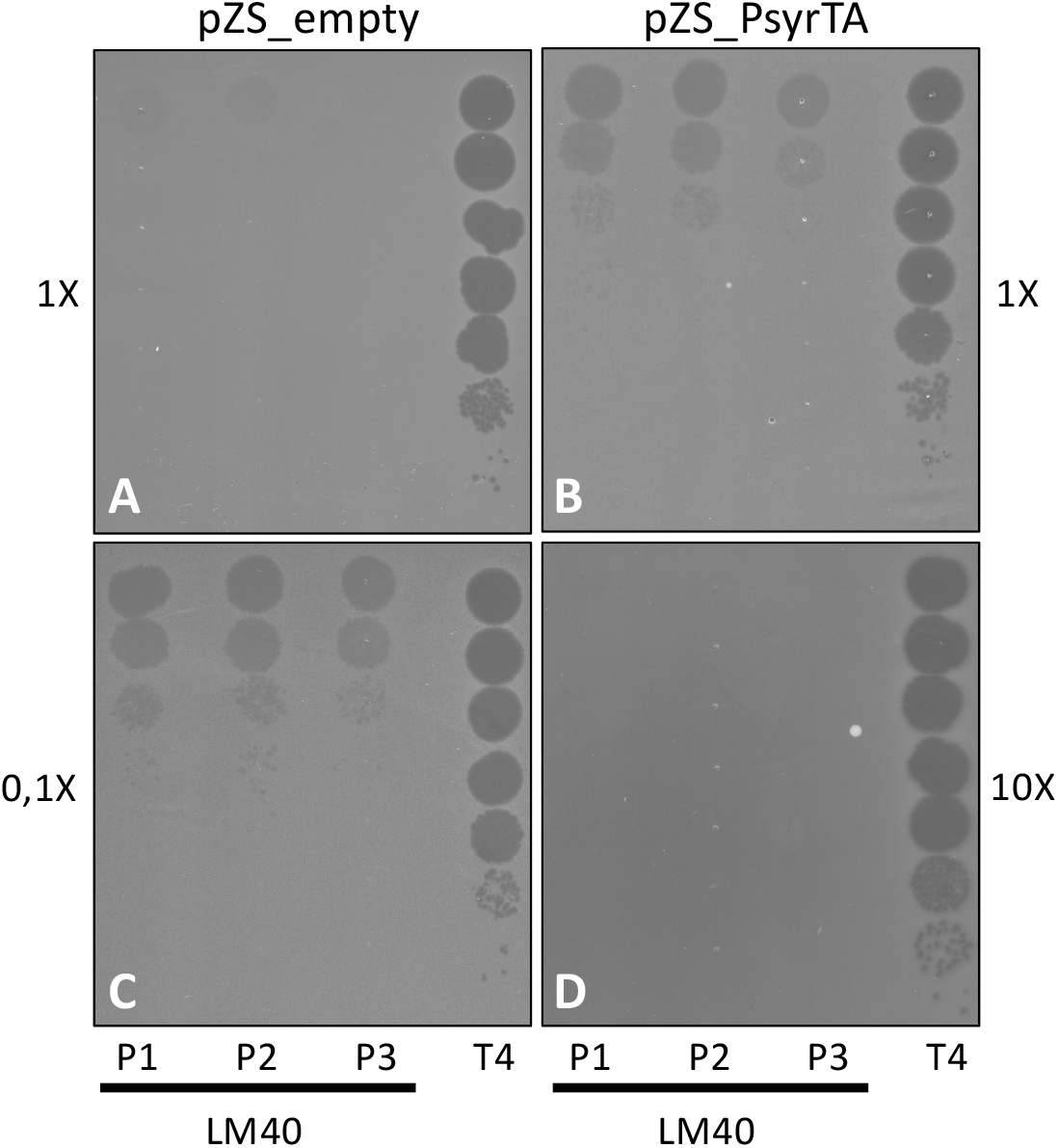
Impact of bacterial lawn density on LM40_P phage infectivity in the K-12 MG1655 strain expressing PsyrTA and its parental strain. A, B. The bacterial lawn was inoculated directly from cultures at OD600 = 0.6 (1x). C, D. The parental and PsyrTA strains were inoculated at dilutions of 1/10 or 10x, respectively, from cultures at OD600 = 0.6. For each infectivity test, 3 μL of 10-fold serially diluted phage was spotted. The T4 phage was used as a positive control for lysis.

**Supplementary Table 1**. List of phages infecting NILS69

**Supplementary Table 2**. List of predicted defence systems and toxin antitoxin systems encoded in NILS69

**Supplementary Table 3**. Lists of plasmids, strains and primers used in this study

