## Supplementary data for "Systematic functional assessment of anti-phage systems in their native host"

**Electronic Supplementary material**

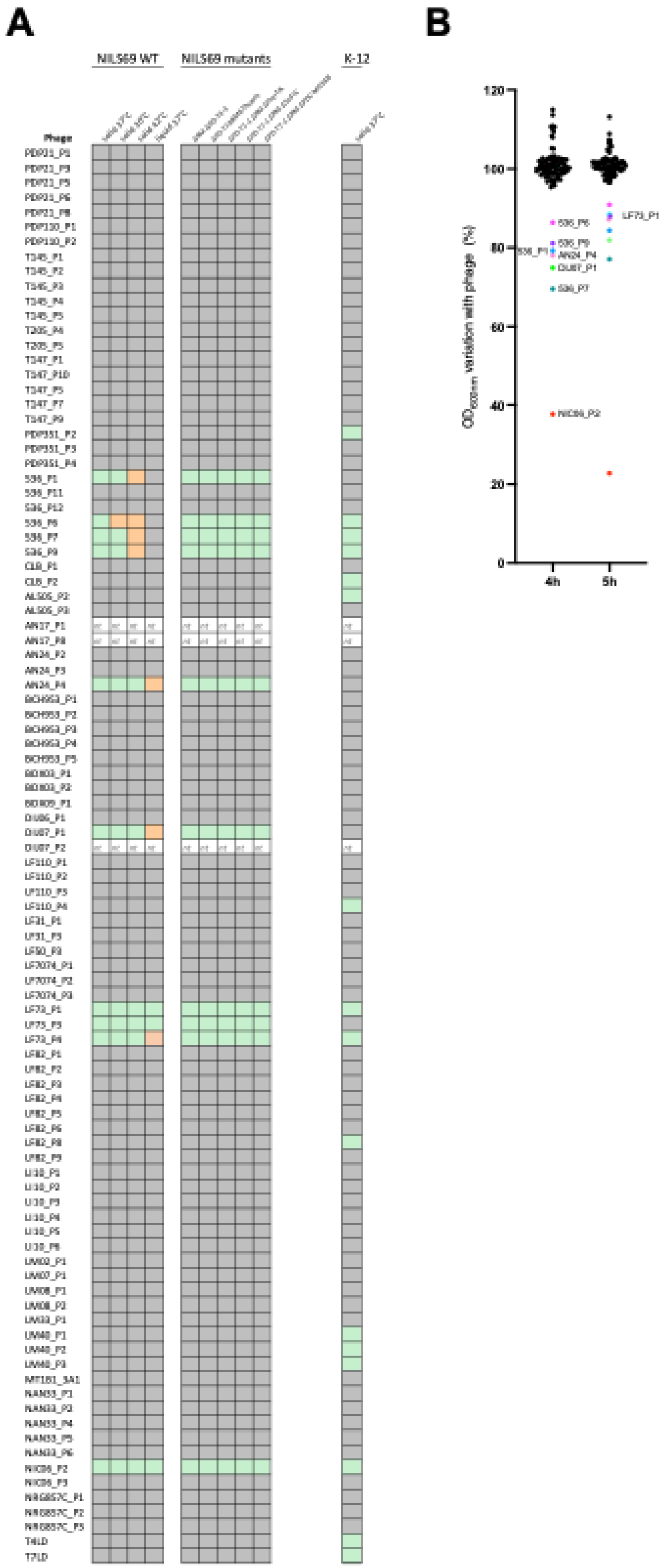


**Supplementary Figure 1. Screening of the Antonina Guelin phage collection on NILS69 and K-12 in solid (A) and liquid (B) infection assays.**
“nt” stands for “not tested”, the corresponding phages could not be amplified on their production strain. Green and orange squares indicate detectable phage infection (i.e. plaques). Orange square indicate cases where only faint lysis could be observed. Grey indicates no lysis

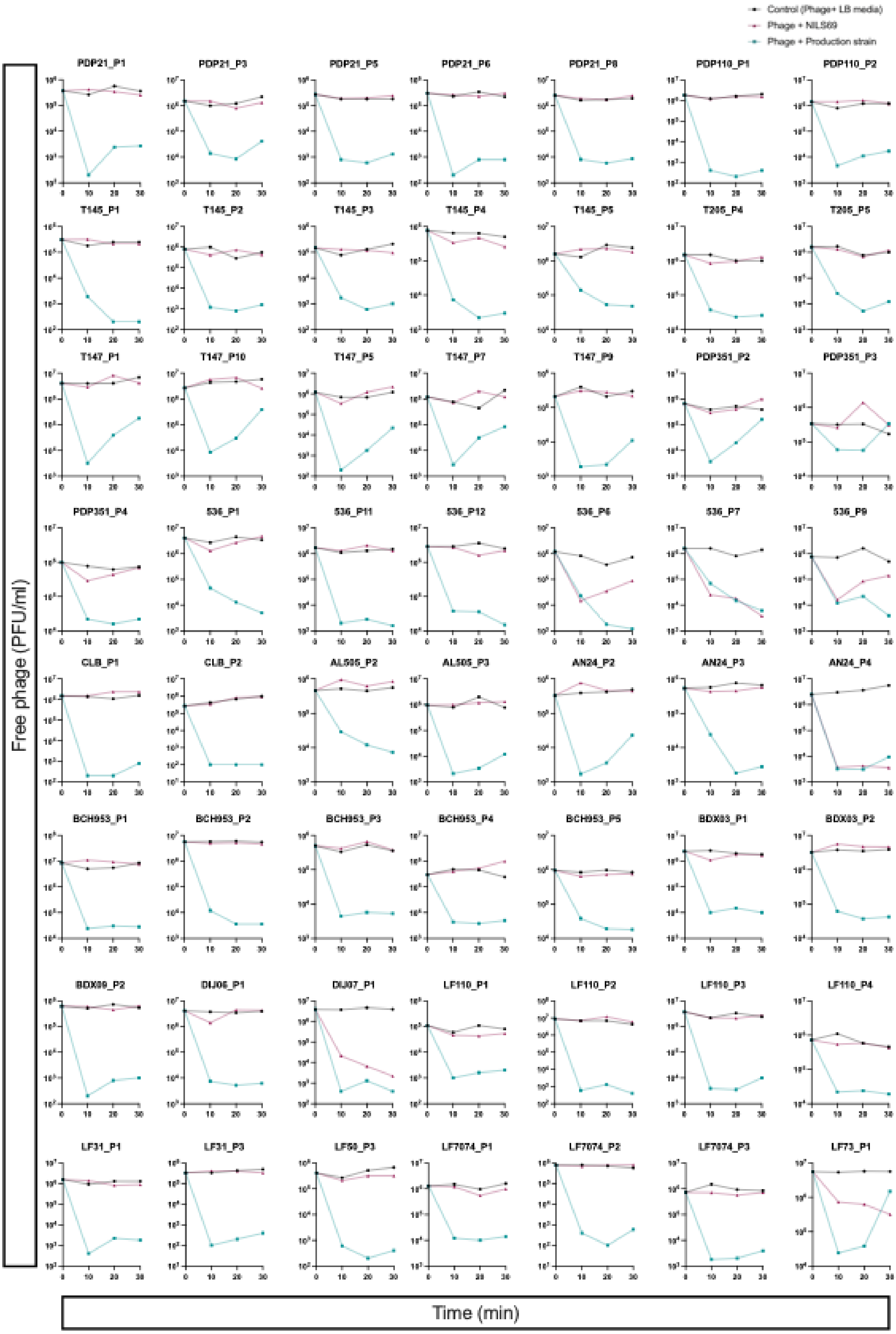
Control (Phage+ LB media)
Phage + NILS69
Phage + Production strain
PDP21_P1
PDP21_P3
PDP21_P5
PDP21_P6
PDP21_P8
PDP110_P1
PDP110_P2
T145_P1
T145_P2
T145_P3
T145_P4
T145_P5
T205_P4
T205_P5
T147_P1
T147_P10
T147_P5
T147_P7
T147_P9
PDP351_P2
PDP351_P3
PDP351_P4
536_P1
536_P11
536_P12
536_P6
536_P7
536_P9
CLB_P1
CLB_P2
AL505_P2
AL505_P3
AN24_P2
AN24_P3
AN24_P4
BCH953_P1
BCH953_P2
BCH953_P3
BCH953_P4
BCH953_P5
BDX03_P1
BDX03_P2
BDX09_P2
DIJ06_P1
DIJ07_P1
LF110_P1
LF110_P2
LF110_P3
LF110_P4
LF31_P1
LF31_P3
LF50_P3
LF7074_P1
LF7074_P2
LF7074_P3
LF73_P1
Free phage (PFU/ml)
Time (min)

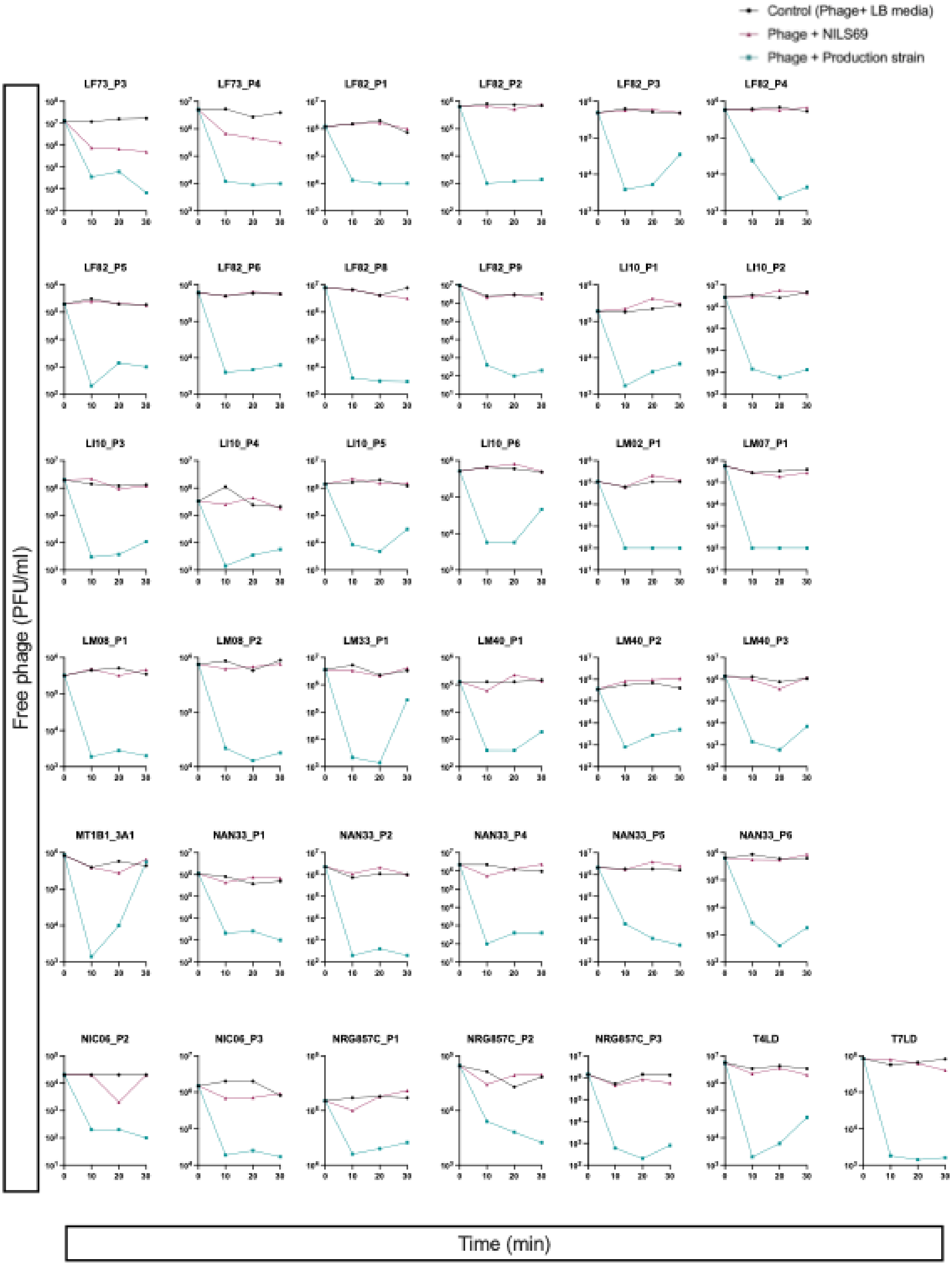


**Supplementary Figure 2. Adorption assays of 93 phages from the Antonina Guelin on NILS69 and on their control strains (phage production strains)**

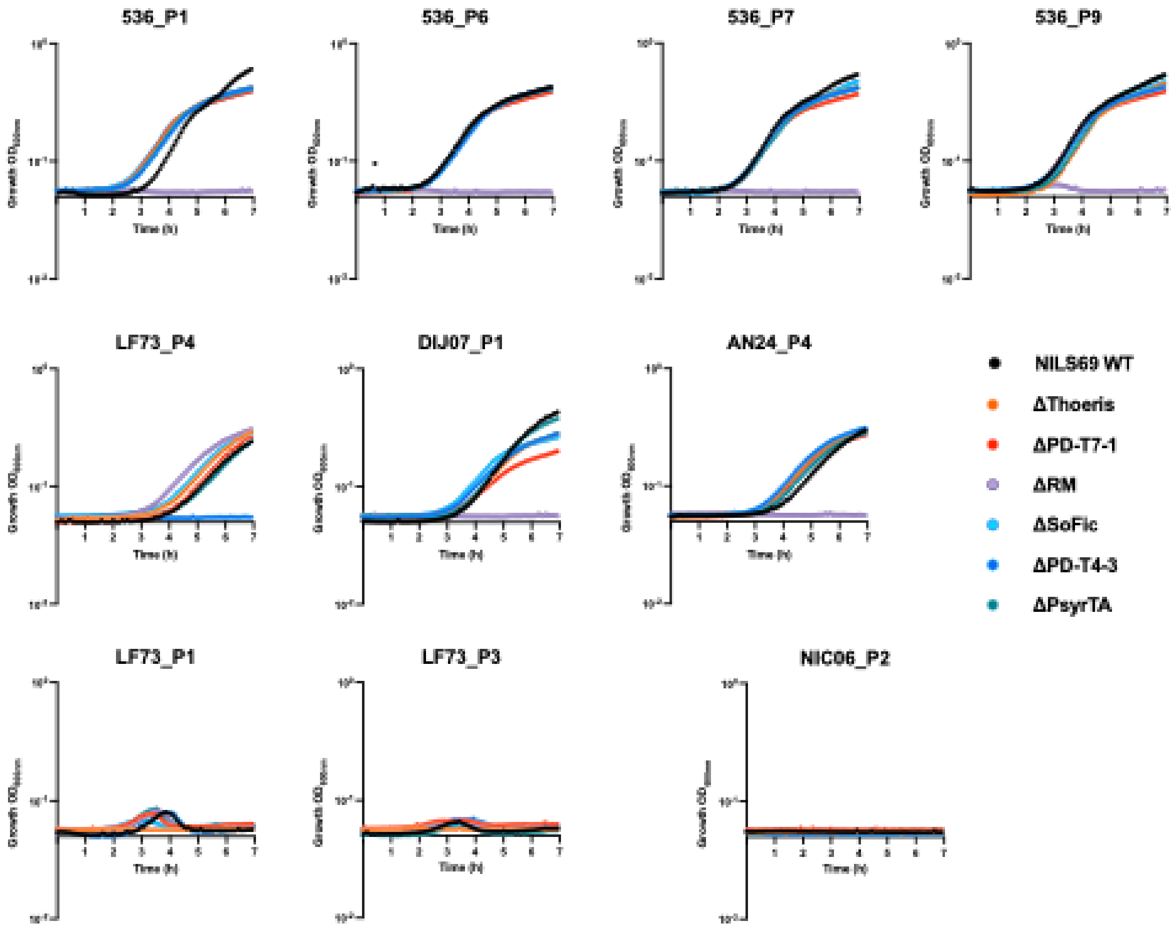


**Supplementary Figure 3. Growth kinetic of NILS69 and mutant dertivative in the presence of the infective phages at a MOI of 1**

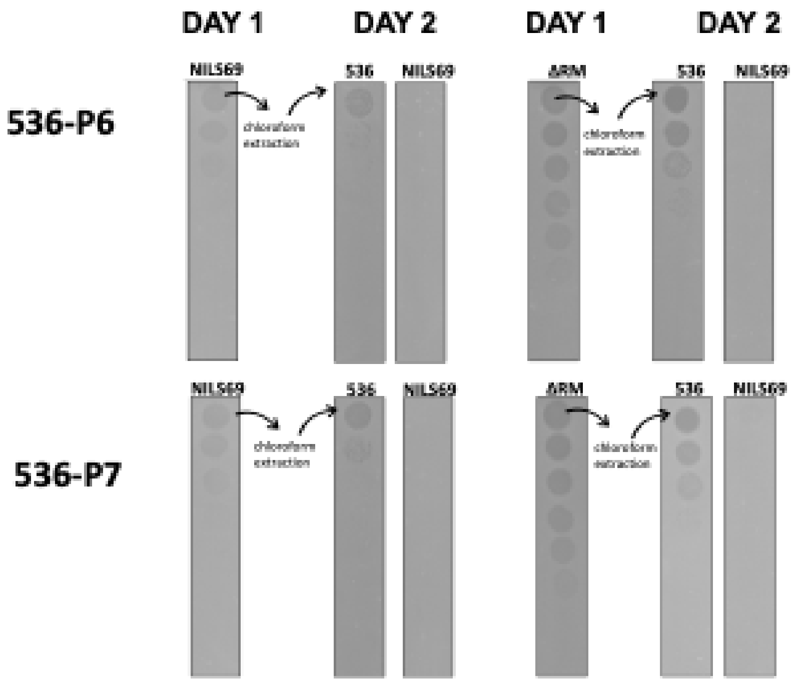


**Supplementary Figure 4. Assessment of escape phage mutants**

Phages 536_P6 and 536_P7 lysing NILS69 or NILS69ΔRM were harvested from the plate and purified through chloroform extraction (Day 1). These phages were subjected to 10-fold serial dilutions and spotted on lawn of strain 536, NILS69 or NILS69ΔRM (Day 2).

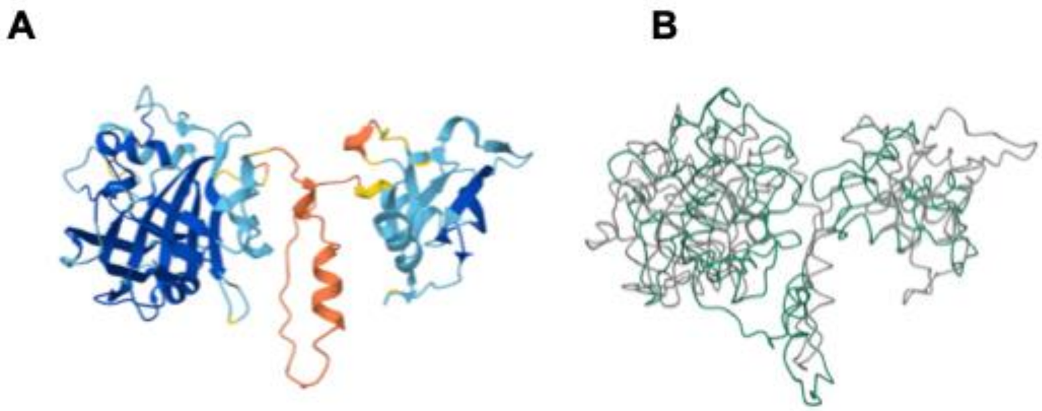


**Supplementary Figure 5: Predicted structure of HNH endonuclease 3**

A. HNH endonuclease 3 encoded in NILS69 has 100% identity with the amino acid sequence of a HNH endonuclease (Uniprot A0A3K0QCZ9), which has a predicted structure in AlphaFold (average pLDDT score of 80,56)

B. Structural alignment of NILS69_HNH endonuclease and K-12 MG1655_EcoKMrcA (e-value 2.48e-4; sequence identity 20.6%)

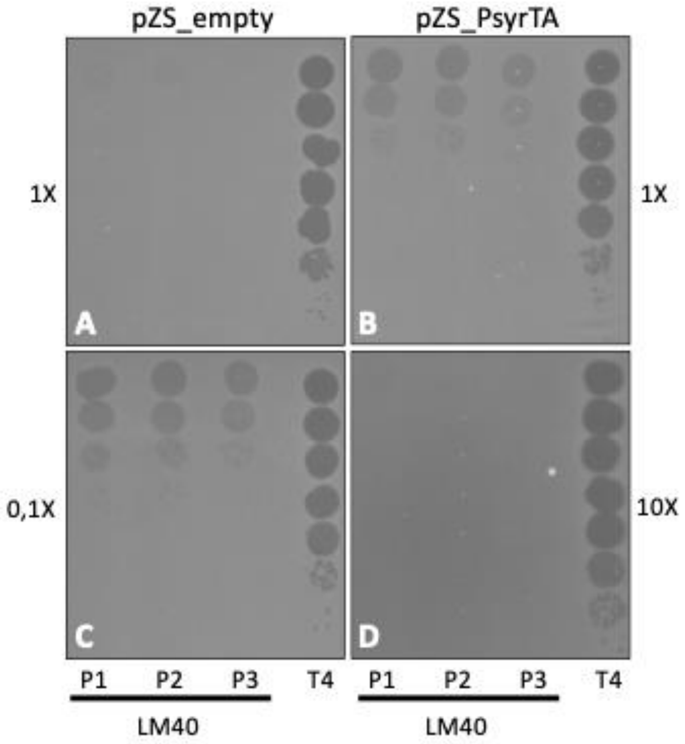


**Supplementary Figure 6: Impact of bacterial lawn density on LM40_P phage infectivity in the K-12 MG1655 strain expressing PsyrTA and its parental strain.**
A, B. The bacterial lawn was inoculated directly from cultures at OD600 = 0.6 (1x).
C, D. The parental and PsyrTA strains were inoculated at dilutions of 1/10 or 10x, respectively, from cultures at OD600 = 0.6. For each infectivity test, 3 µL of 10-fold serially diluted phage was spotted. The T4 phage was used as a positive control for lysis.

**Supplementary Table 1.** List of phages infecting NILS69

**Supplementary Table 2.** List of predicted defence systems and toxin antitoxin systems encoded in NILS69

**Supplementary Table 3.** Lists of plasmids, strains and primers used in this study
